# *TaAPO-A1*, an ortholog of rice *ABERRANT PANICLE ORGANIZATION 1*, is associated with total spikelet number per spike in elite hexaploid winter wheat varieties (*Triticum aestivum* L.)

**DOI:** 10.1101/659813

**Authors:** Quddoos H. Muqaddasi, Jonathan Brassac, Ravi Koppolu, Jörg Plieske, Martin W. Ganal, Marion S. Röder

## Abstract

We dissected the genetic basis of total spikelet number (TSN) along with other traits, namely spike length (SL) and flowering time (FT) in a panel of 518 elite European winter wheat varieties. Genome-wide association studies based on 39,908 SNP markers revealed highly significant quantitative trait loci (QTL) for TSN on chromosomes 2D, 7A, and 7B, for SL on 5A, and FT on 2D, with 2D-QTL being the functional marker for the gene *Ppd-D1*. The physical region of the 7A-QTL for TSN revealed the presence of an ortholog to *APO1* – a rice gene that positively controls spikelet number on panicles. Interspecific analyses of *TaAPO-A1* orthologs showed that it is a highly conserved gene important for floral development, and present in a wide range of terrestrial plants. Intraspecific studies of the wheat ortholog *TaAPO-A1* across wheat genotypes revealed a polymorphism in the highly conserved F-box domain, defining two haplotypes. A KASP maker developed on the polymorphic site showed a highly significant association of *TaAPO-A1* with TSN, explaining 23.2% of the genotypic variance. Also, the *TaAPO-A1* alleles showed weak but significant differences for SL and grain yield. Our results demonstrate the importance of wheat sequence resources to identify candidate genes for important traits based on genetic analyses.

## Introduction

The wheat spike and its architecture are key components for improving grain yield. In the recent past, several genes controlling spike morphology have been investigated and described in temperate cereals (Gauley and Boden 2019; Koppolu and Schnurbusch 2019). From a plant breeder’s viewpoint, most spike morphological traits in wheat such as spike length and spikelet number behave as quantitative traits, and various QTL and association studies have recently been published (Deng et al. 2017; Guo et al. 2017; Liu et al. 2018; Sakuma et al. 2019; Würschum et al. 2018; Zhai et al. 2016). High associations and prediction abilities for total and fertile spikelet number as well as spike length and grain yield were also reported (Guo et al. 2018).

Nevertheless, only a few cloned genes for the trait number of spikelet pairs in wheat are available, among them is the *Q*-gene which played a major role in wheat domestication and encodes an *AP2* transcription factor (Faris et al. 2003). The domesticated allele *Q* confers a free-threshing character, a sub-compact spike (Greenwood et al. 2017), and is regulated by microRNA172 (Debernardi et al. 2017). Also, genes related to heading date are involved in spikelet meristem identity determination. For example, the photoperiodism gene *Ppd* was reported to influence spikelet primordia initiation (Ochagavía et al. 2018). Mutants of the *FLOWERING LOCUS T2* (*FT2*) in wheat showed a significant increase in the number of spikelets per spike with an extended spike development period accompanied by delayed heading time (Shaw et al. 2018). Moreover, *Ppd-1* and *FT* were reported as regulators of paired spikelet formation resulting in increased number of grain producing spikelets (Boden et al. 2015). Mutants of the MADS-box genes, e.g., *VRN1* or *FUL2* showed increased number of spikelets per spike, likely due to a delayed formation of the terminal spikelet (Li et al. 2019) and a putative ortholog to rice *MOC1* regulating axillary meristem initiation and outgrowth was associated with spikelet number per spike in wheat (Zhang et al. 2015).

The *ABERRANT PANICLE ORGANIZATION 1* (*APO1*) gene in rice was reported to affect the inflorescence structure severely (Ikeda et al. 2005). It encodes an F-box protein which is an ortholog of *UNUSUAL FLORAL ORGAN* (*UFO*), regulating floral identity in *Arabidopsis* (Samach et al. 1999; Wilkinson and Haughn 1995). Characterization of rice *apo1* mutants revealed that *APO1* positively controls spikelet number by suppressing the precocious conversion of inflorescence meristems to spikelet meristems. Besides this, *APO1* was associated with the regulation of the plastochron, floral organ identity, and floral determinacy (Ikeda et al. 2007). Four dominant mutants of *APO1* with elevated expression levels of *APO1* produced increased number of spikelets by a delay in the programmed shift to spikelet formation. Ectopic overexpression of *APO1* resulted in increased meristem size caused by different rates of cell proliferation. It was concluded that the level of *APO1* activity regulates the inflorescence form through the control of meristematic cell proliferation (Ikeda-Kawakatsu et al. 2009).

In the present study, we investigated the inheritance and genetic basis of total spikelet number (TSN) per spike, spike length and flowering time as component traits of grain yield in an elite European winter wheat panel. Our findings show the complex genetic architecture of the investigated traits, and that *TaAPO-A1* – an ortholog of rice *APO1*, which is vital for inflorescence development, is associated with TSN determination in wheat. Intraspecific sequence analyses of *TaAPO-A1* revealed that polymorphisms were forming distinct haplotypes while intraspecific studies showed the conserved nature of this gene across terrestrial plant species.

## Materials and methods

### Phenotypic data analyses

The data for total spikelet number (TSN), spike length (SL), and flowering time (FT) were collected on an elite European winter wheat panel comprising of 518 varieties. The whole panel was grown in the experimental fields of Leibniz Institute of Plant Genetics and Crop Plant Research (IPK) Gatersleben, Germany in plots of 2 m^2^ as single replication in three cropping seasons (2015/16; 2016/17; and 2017/18), henceforth called environments. The traits TSN and SL were recorded in two environments (2016/17 and 2017/18) from ten spikes per plot as the total number of spikelets and spike length in centimeters (cm) from basal spikelet to the top of a spike by excluding the awns. The arithmetic mean of TSN and SL from ten spikes were calculated to represent the genetic value of traits in the individual environments. Flowering time was recorded in all three environments by counting the number of days from the first of January to when approximately half of the spikes in a plot flowered. The phenotypic data for grain yield estimated in eight environments were taken from the previous study for comparison purposes (Schulthess et al. 2017). A linear mixed-effect model was used for across environment phenotypic data analysis as:

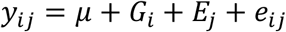

where, *y*_*ik*_ is the phenotypic record of the *i*^*th*^ genotype in the *j*^*th*^ environment, μ is the common intercept term, *G*_*i*_ is the effect of the *i*^*th*^ genotype, *E*_*j*_ is the effect of the *j*^*th*^ environment, and *e*_*ij*_ denotes the corresponding error term. All effects, except the intercept, were assumed random to calculate the individual variance components. The broad-sense heritability (*H*^2^) was calculated as:

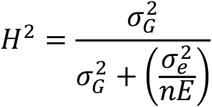

where, 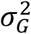 and 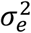 denote the variance components of the genotype and the error, respectively; and *nE* denotes the number of environments. To calculate the best linear unbiased estimations (BLUEs), the intercept and the genotypic effects were assumed fixed in the above model.

### Genotypic data analyses, population structure, and linkage disequilibrium

All 518 varieties were extensively genotyped with the 35k Affymetrix and 90k iSELECT single nucleotide polymorphism (SNP) arrays (Allen et al. 2017; Wang et al. 2014) which generated in total 116,730 SNP markers (35k = 35,143; 90k = 81,587). We also genotyped the whole panel with functional markers for the candidate genes such as photoperiodism (*Ppd-D1*), reduced height (*Rht*), and vernalization (*Vrn1*). The quality of the marker data was improved by removing the markers harboring >10% heterozygous or missing calls and markers with a minor allele frequency of <0.05. The mean of both alleles imputed the remaining missing data. The quality control resulted in a total of 39,908 markers, which were used in subsequent analyses.

Population structure based on marker genotypes was examined by principal component (PC) analysis. The first two PCs were drawn to see the clustering among varieties. Moreover, the genetic relatedness among varieties was evaluated by an additive variance-covariance genomic relationship matrix. To infer the hidden population sub-structuring, an inference algorithm LEA (Landscape and Ecological Association Studies) was used by assuming ten ancestral populations (*K* = 1-10). The function *snmf*, which provides the least squares estimates of ancestry proportions and estimates an entropy criterion to evaluate the quality fit of the model by cross-validation, was used. The number of ancestral populations best explaining the data can be chosen by using the entropy criterion. We performed ten repetitions for each *K*, and the optimal repetition demonstrating the minimum cross-entropy value was used to visualize clustering among varieties via bar plots (Frichot and François 2015).

Linkage disequilibrium (LD), the non-random association of alleles at different loci, was measured as the squared correlation (*r*^2^) among markers. The genetic mapping positions of the markers for both arrays were adopted from the data generated for the International Triticeae Mapping Initiative (ITMI) DH population, as described in Sorrells et al. (2011). Although inter and intra-chromosomal LD among the loci varies, genome-wide calculation of LD gives a global estimate about the genetic map distance over which LD decays in the given population. The genome-wide (global) LD was calculated only from the mapped markers.

### Genome-wide association studies

Genome-wide association studies (GWAS) were performed on data taken from the individual environment and SNPs passing the quality criteria *plus* the functional gene markers. Let *n* be the number of varieties and *p* be the predictor marker genotypes. A standard linear mixed-effect model following Yu et al. (2006) was used to perform GWAS as:

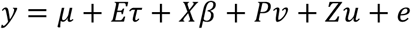

where, *y* is the *n* × 1 vector of phenotypic record of each genotype in each environment, μ is the common intercept, *τ*, *β*, *ν*, *u*, and *e* are the vectors of the environment, marker, population (principal components), polygenic background, and the error effects, respectively; *E*, *X*, *P*, and *Z* are the corresponding design matrices. In the model, *μ, τ, β*, and *ν* were assumed fixed while *u* and *e* as random with 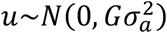 and 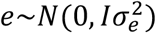. The *n* × *n* variance-covariance additive relationship matrix (*G*) was calculated from *n* × *p* matrix *W* = (*w*_*ik*_) of marker genotypes (being 0, 1 or 2) as 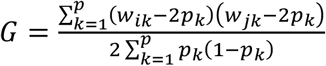 where, *w*_*ik*_ and *w*_*jk*_ are the profiles of the *k*^*th*^ marker for the *i*^*th*^ and *j*^*th*^ variety, respectively; *p*_*k*_ is the estimated frequency of one allele in *k*^*th*^ marker, as described by VanRaden (2008).

As population stratification and familial relatedness can severely impact the power to detect true marker-trait association (MTA) in GWAS, different statistical models were used to avoid spurious MTA viz., (1) general linear model (*naive*), (2) population structure correction via principal components (*PCs*), (3) correction of familial relatedness via genomic relationship matrix (*G*), and (4) correction of population structure and relatedness via *PCs* and *G*. It is expected that using both *PCs* and *G*, in the model can enhance the accuracy of GWAS. Along with this, environmental fixed effects were assigned in all model scenarios. The models described above were compared by plotting expected versus the observed −log_10_(*P* − value) in a quantile-quantile plot and the best model was determined by checking how well the observed −log_10_(*P* − value) aligned with the expected.

To declare the presence of MTA, a false discovery rate (FDR) <0.05 to account for multiple testing was applied (Benjamini and Hochberg 1995). Following Utz et al. (2000), the percentage of total genotypic variance (*p*_*G*_) explained by all the QTL passing the FDR threshold was determined as 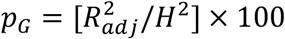 where, 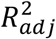 was calculated by fitting all the MTA in a multiple linear regression model in the order of ascending *P*-values and *H*^2^ is the broad-sense heritability. The *p*_*G*_ values of individual QTL were accordingly derived from the sum of squares of the QTL (SS_QTL_) in the linear model.

### Candidate gene identification, haplotype analysis by exploiting resources from The 10+ Wheat Genome Project, and KASP marker development

We narrowed-down the QTL region, and BLASTed sequences of all the significant markers present within the genetically defined region onto the physical map of the corresponding chromosome of the reference sequence of the wheat genome which yielded significant physical region (Altschul et al. 1990; Consortium 2018). Afterward, the gene identifiers (gene-IDs) present within the physical region and their annotated functional descriptions were retrieved. Among them was a most likely candidate gene *TaAPO-A1* for TSN.

*The 10+ Wheat Genome Project* is an international collaborative effort that aims to assemble the genomes of more than ten wheat varieties bred in different countries to characterize the wheat pan-genome (http://www.10wheatgenomes.com/). We retrieved the genomic sequence of *TaAPO- A1* for ten wheat varieties from *The 10+ Wheat Genome Project* and aligned the sequences to observe the haplotype structures. The SNP that revealed a clear haplotype structure was used to design a Kompetitive Allele Specific PCR (KASP) marker in the candidate gene. The allele-wise phenotypic distribution of the investigated traits with the gene-specific KASP marker was analyzed by plotting the boxplots. The significance (*P-*values) between the mean values of genotypes harboring different KASP marker alleles was determined by two-sided t-test. Moreover, we performed a second round of GWAS by incorporating the gene-specific KASP marker in the original SNP matrix to determine whether it associates with the phenotypes. The GWAS parameters were kept the same as described above.

### Multiple sequence alignment and phylogenetic analyses

The TaAPO-A1 protein sequence (corresponding to *TraesCS7A01G481600*) was used as a BLAST query to retrieve the monocot, dicot and Bryophyte orthologs from EnsemblPlants (http://plants.ensembl.org/index.html) and Phytozome v12.1 (https://phytozome.jgi.doe.gov/pz/portal.html) databases. The orthologous protein sequences were aligned using ClustalW in Geneious v.11.0.5 (Kearse et al. 2012). The protein alignment was used to infer a maximum likelihood (ML) phylogeny. The JTT matrix (Jones et al. 1992) was identified as the best-fitting model of protein evolution with ProtTest 3 (Darriba et al. 2011; Guindon and Gascuel 2003) and the Akaike Information Criterion (AIC). The evolutionary history among *TaAPO-A1* orthologs across various plant species was inferred using RAxML v8.2.12 (Stamatakis 2014) with PROTGAMMAJTT model, rapid bootstrapping of 100 replicates, and search for best-scoring ML tree (options “-f a -x 1 -# 100”). The consensus tree was further processed to collapse branches with bootstrap support lower than 50%, and the tree was rooted with the Bryophytes *Physcomitrella patens* and *Selaginella moellendorffii* as outgroup.

## Results

### Total spikelet number per spike is significantly correlated with spike length, flowering time, and grain yield

The assessment of total spikelet number (TSN) per spike, spike length (SL), and flowering time (FT) were performed in the field trials on 518 elite European winter wheat varieties (including 15 spring type wheat varieties as an outgroup). The trait grain yield (GY) was assessed in multiple environment field trials on a subset (in total, 372) of varieties in a previous study (Schulthess et al. 2017). The best linear unbiased estimations (BLUEs) of all traits approximated normal distribution and showed wide variation (Fig. 1a-d; Table S1). The ANOVA showed that genotypic 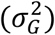 and environmental 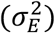 variation was significantly (*P* < 0.001) larger than zero (Table 1). The broad-sense heritability ranged from 0.68 to 0.89 which indicates the good quality of the phenotypic data and its potential for use in genome-wide association (GWAS) studies to map the quantitative trait loci (QTL) underlying the traits (Table 1). We analyzed the Pearson product moment correlation (*r*) among the BLUEs of investigated traits, which revealed that TSN was positively and significantly correlated with SL, FT, and GY (Fig. 1e). The TSN and SL showed the highest correlation among the analyzed traits (*r* = 0.46; *P* < 0.001) whereas SL showed almost a null correlation with FT and GY suggesting that FT augments GY mainly by influencing TSN in wheat.

**Table 1.** Summary statistics of the investigated traits, namely total spikelet number (TSN), spike length (SL), flowering time (FT), and grain yield (GY).

| Parameter | TSN | SL | FT | GY |
| --- | --- | --- | --- | --- |
| Minimum | 14.38 | 7.90 | 144.96 | 73.94 |
| Mean | 19.45 | 10.12 | 153.38 | 96.74 |
| Maximum | 24.75 | 13.05 | 159.61 | 110.71 |
| $\sigma_G^2$ | 1.71<sup>a</sup> | 0.50<sup>a</sup> | 3.42<sup>a</sup> | 22.89<sup>a</sup> |
| $\sigma_E^2$ | 1.75<sup>a</sup> | 1.63<sup>a</sup> | 6.30<sup>a</sup> | 94.51<sup>a</sup> |
| $\sigma_e^2$ | 1.60 | 0.44 | 1.90 | 23.74 |
| $H^2$ | 0.68 | 0.70 | 0.84 | 0.89 |
| $nE$ | 2 | 2 | 3 | 8 |

$\sigma_G^2$ = genotypic variance; $\sigma_E^2$ = environmental variance; $\sigma_e^2$ = residual variance; $H^2$ = broad-sense heritability; $nE$ = number of environments; a = significant at <0.001 probability level.

**Figure 1:**
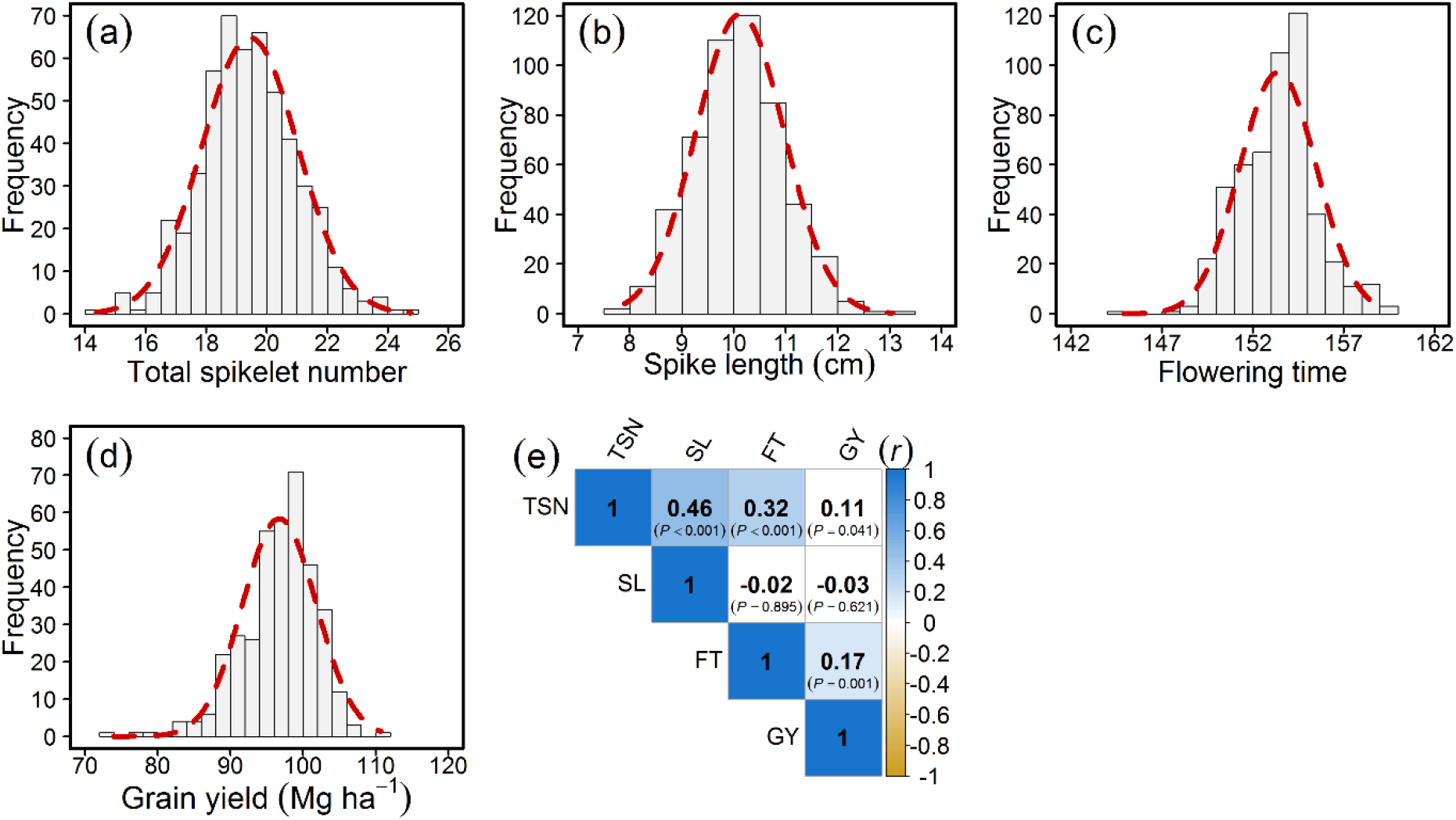
Distribution and correlation of the investigated traits in a panel of 518 elite European winter wheat varieties. Distribution of (a) Total spikelet number (TSN), (b) Spike length (SL), (c) Flowering time (FT), and (d) Grain yield (GY); (e) Pearson product moment correlation (*r*) among the investigated traits. *P*-value denotes the significance of the respective correlation.

### High-density marker arrays reveal the absence of distinct sub-populations and sharp LD decay

The whole wheat panel was extensively genotyped with high-density SNP arrays and functional markers for the genes *Ppd-D1*, *Rht-B1*, *Rht-D1*, *Vrn-A1*, *Vrn-B1*, and *Vrn-D1*, which resulted in 39,908 high-quality markers. The population structure analyzed with marker genotypes by PC analysis resulted in the absence of distinct sub-populations with the first two PCs representing only 11.3% of the variation (Fig. 2). The high familial relatedness and non-existence of distinct sub-populations were further supported by plotting a heat map of the genomic relationships among the wheat varieties (Fig. S1) and by the structure-like inference algorithm LEA, which resulted in the sub-populations being distinguished but with a slight entropy shift. The bar plots indicated admixed and weak sub-populations (Fig. S2).

**Figure 2:**
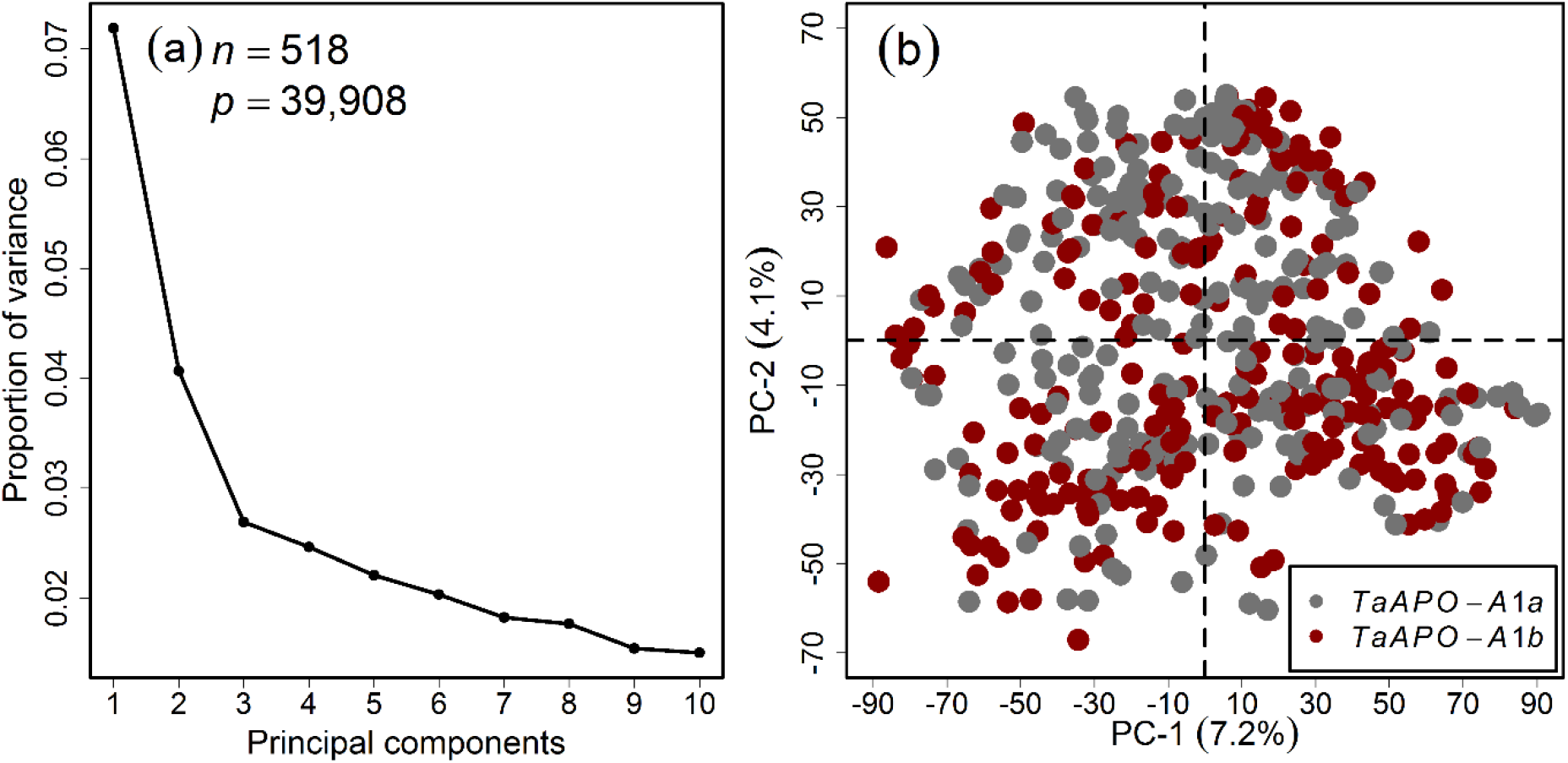
Principal component (PC) analysis on the wheat marker loci combined from the 35k and 90k single nucleotide polymorphism arrays. (a) Scree plot showing the first ten PCs and their corresponding proportion of variance, (b) Scatterplot showing the absence of pronounced clustering among the varieties. Different colors represent the *TaAPO-A1* alleles. *n* and *p* denote the number of varieties and the marker genotypes used in the analysis, respectively.

Linkage disequilibrium (LD) between the marker genotypes determines the number of markers needed to perform GWAS. Genome-wide LD was performed with the mapped marker genotypes which resulted in rapid LD decay with increasing the genetic map (cM) distances, with first and third quantile dropping to 0.002 and 0.028, respectively; and the mean and median values equaling 0.051 and 0.008, respectively (Fig. 3a). The sub-genome-wise distribution of the markers varied, with the highest markers mapping on B-genome, followed by A- and D-genomes (Fig. 3b). Although the whole panel was genotyped with state-of-the-art genotyping arrays, the sub-genome-wise distribution of marker genotypes suggests that marker density could be improved especially for D-genome.

**Figure 3:**
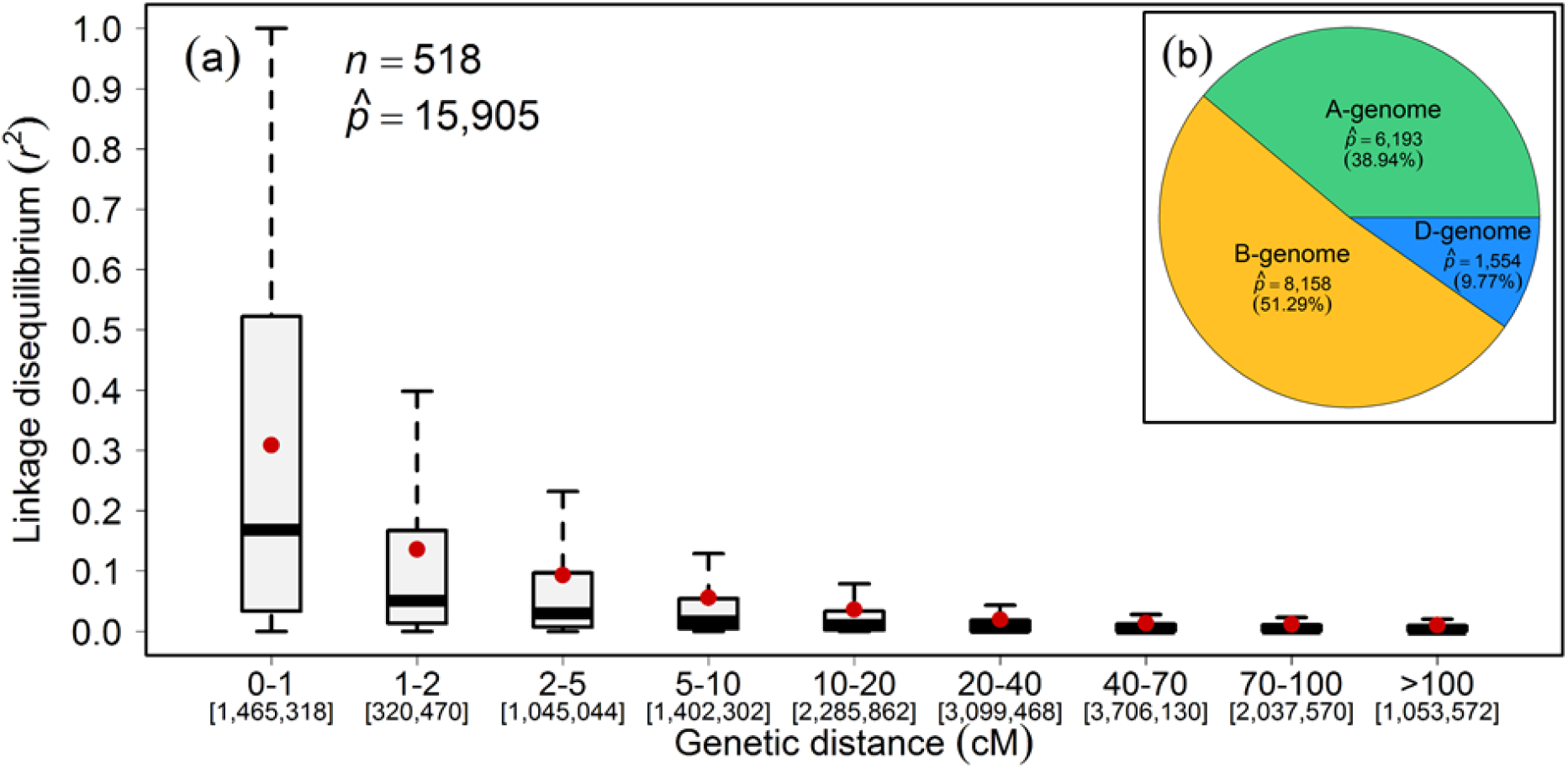
Genome-wide decay of linkage disequilibrium (LD; *r*^2^) as a function of genetic map distance (cM) between the marker loci in the population of European winter wheat varieties. (a) Boxplots represent the LD-decay, (b) Sub-genome-wise distribution of mapped marker loci. Red dots within the boxplots represent the mean. *n* and 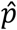 denote the number of varieties and mapped marker loci, respectively.

### GWAS identifies large-effect QTL for TSN on chromosome 7A in wheat varieties

Among the different GWAS models used in our study, we observed that the *PC*_[1-3]_+*G* model could best control the spurious MTA. Our GWAS analyses identified QTL on chromosomes 2D, 7A, and 7B for TSN (Fig. 4a-b; Table S1a), for SL on chromosome 5A (Fig. S3, Table S1b), and for FT on chromosome 2D (Fig. S4; Table S1c). The QTL on chromosome 2D identified for TSN and FT was very likely the gene *Ppd-D1*. Of particular interest is the photoperiod insensitive allele *Ppd-D1a* that significantly reduced the TSN (Fig. 4a, f). The phenotypic data for GY were analyzed to investigate if there exists any significant correlation between the identified marker alleles and GY. The total proportion of genotypic variance (*p*_*G*_) imparted by the identified QTL amounted to 65.44% for TSN, 15.15% for SL, and 31.58% for FT. A relatively low *p*_*G*_ for SL and FT is the result of the identification of only one mapped MTA for each trait.

**Figure 4:**
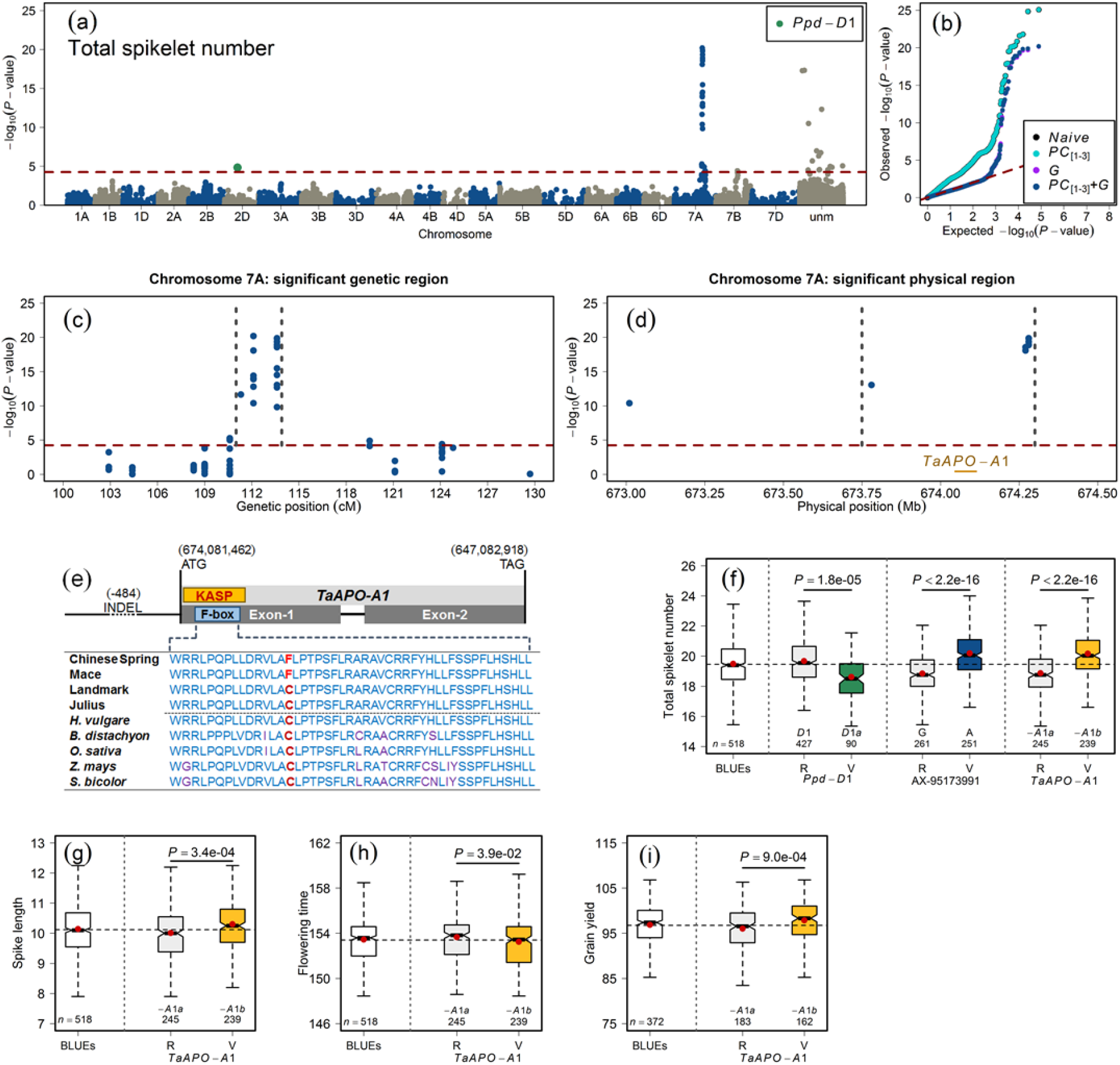
Summary of genome-wide association studies of total spikelet number per spike in the population of 518 European winter wheat varieties. (a) Manhattan plot shows the distribution of marker significance −log_10_(*P* − value) along the chromosomes. The correction for population stratification and familial relatedness was performed by using the first three principal components (*PC*_[1-3]_) and an additive genomic relationship matrix (*G*) in a linear mixed-effect model. The red dashed line marks the multiple testing criteria of false discovery rate (FDR) <0.05, **(b) Quantile-quantile plot showing the distribution of observed versus expected (red dashed line) −log_10_(*P* − value).** The general linear model (*naive*) without correction for population structure, the *PC*_[1-3]_ model (population structure corrected with the first three *PCs*), the *G* model (familial relatedness corrected with a genomic relationship matrix), and the *PC*_[1-3]_+*G* model (population structure and familial relatedness corrected with *PCs* and the *G* matrix). The color code for different models is given in the figure legend, **(c) Significant genetic region on chromosome 7A for TSN in wheat.** The gray vertical dashed lines mark the highly significant genetic region, **(d) Significant physical region on chromosome 7A for TSN in wheat.** The gray vertical dashed lines mark the highly significant physical region, **(e) Gene structure of *TaAPO-A1*.** The orange box represents the location of the KASP marker developed to exploit the variation in the F-box domain (highlighted in blue color). The horizontal line before the first exon depicts promotor region harboring INDEL and corresponding position. The first four rows represent the F-box sequences of wheat varieties (courtesy: *The 10+ Wheat Genomes Project*) and the second four rows represent the F-box domain of closely related species viz., *Hordeum vulgare*, *Brachypodium distacyon*, *Oryza sativa*, *Zea mays*, and *Sorghum bicolor*. The non-synonymous mutation is highlighted in red color. The location of start and stop codons on chromosome 7A are given in the figure, **(f) Allele-wise phenotypic distribution of the most significant markers and KASP marker for *TaAPO-A1* associated with (f) TSN, (g) Spike length, (h) Flowering time, and (i) Grain yield.** *P* denotes the significance value of the two-sided t-test used to compare the mean value of marker alleles. In sub-figures (f) to (i), the first boxplots represent the phenotypic distribution of the best linear unbiased estimations (BLUEs) for the respective trait, whereas R and V denote the reference (major) and variant (minor) allele in the investigated population, respectively.

Nevertheless, of interest is the large-effect QTL identified for TSN on chromosome 7A – for which the most significant marker *AX-95173991* is located at 112.10 cM and explained 25.70% of the genotypic variance. This warrants, on the one hand, that the use of 7A-QTL would be beneficial for efficient marker-assisted selection. On the other hand, it made possible the further investigation of 7A-QTL at the physical sequence level to search for candidate genes.

### Significant physical region of chromosome 7A-QTL harbors TaAPO-A1 – a putative candidate gene for TSN in wheat varieties

The significant 7A-QTL region for TSN spanned initially from 110.6 to 124.1 cM (Table S1a). We narrowed down the genetic region with the highly significant MTA with −log_10_(*P* − value) > 10 within 2.3 cM starting from 111.3 to 113.6 cM (Fig. 4c). The alignment of marker sequences present within this most significant genetic region onto chromosome 7A revealed a physical region starting from 673.75 to 674.30 Mb (Fig. 4d) that harbored only ten genes. The functional annotations of these ten genes revealed an interesting candidate gene *TraesCS7A01G481600*; (physical map position: 674,081,462 – 674,082,919 bp) with functional annotation as *Aberrant panicle organization 1* (*APO1*) *protein*. The *APO1* in rice regulates inflorescence architecture and positively controls the total spikelet number by suppressing the precocious conversion of inflorescence meristems to spikelet meristems (Ikeda et al. 2007; Ikeda et al. 2005).

### A KASP marker developed for TaAPO-A1 shows significant association with TSN in wheat varieties

*TaAPO-A1* is a 1,457 bp long gene, and like *APO1* in rice, it has two exons separated by one intron (Fig. 4e). We investigated the variation of *TaAPO-A1* in ten wheat varieties; the sequences were taken from *The 10+ Wheat Genome Project*, which revealed two haplotypes (Fig. 4e; Fig. S6). The first exon harbors a highly conserved F-box domain of 46 amino acid residues across the wheat varieties and other species (Figs. 4e, S6, and S7). Intraspecific sequence analysis of *TaAPO-A1* revealed a non-synonymous mutation in the F-box domain; and out of ten wheat varieties, four (including Chinese Spring) harbored T, while six had G allele. We developed a KASP marker for *TaAPO-A1* harboring this non-synonymous mutation in the F-box domain (Figs. 4e and S6; Table S1a, b). The alleles of the KASP marker were evenly distributed in the variety panel (Figure 2b) and were highly significantly associated with TSN (Fig. 4f; Table S2a). The second round of GWAS was performed by the *TaAPO-A1* KASP marker integrated into the original SNP matrix which further confirmed the significant association of *TaAPO-A1* with TSN, explaining 23.21% of the genotypic variance (Fig. S5; Table S2). The reference allele in the population (represented by *TaAPO-A1a*, with nucleotide G translating to cysteine) was present in 50.62% of the investigated varieties and resulted in an average TSN of 18.83. Whereas the variant allele (represented by *TaAPO-A1b*, with nucleotide T translating to phenylalanine) was present in 49.38% of the varieties and revealed an average TSN of 20.13 (Fig. 4f, Table S1a). The analysis of local linkage disequilibrium performed with the markers present in the 7A-QTL genetic region and the KASP marker for *TaAPO-A1* showed that *TaAPO-A1* was in tight linkage with other markers (Fig. 5). Furthermore, we also observed a rather weak but significant association of the *TaAPO-A1* KASP marker alleles with SL, FT, and GY (Fig. 4g-i).

**Figure 5:**
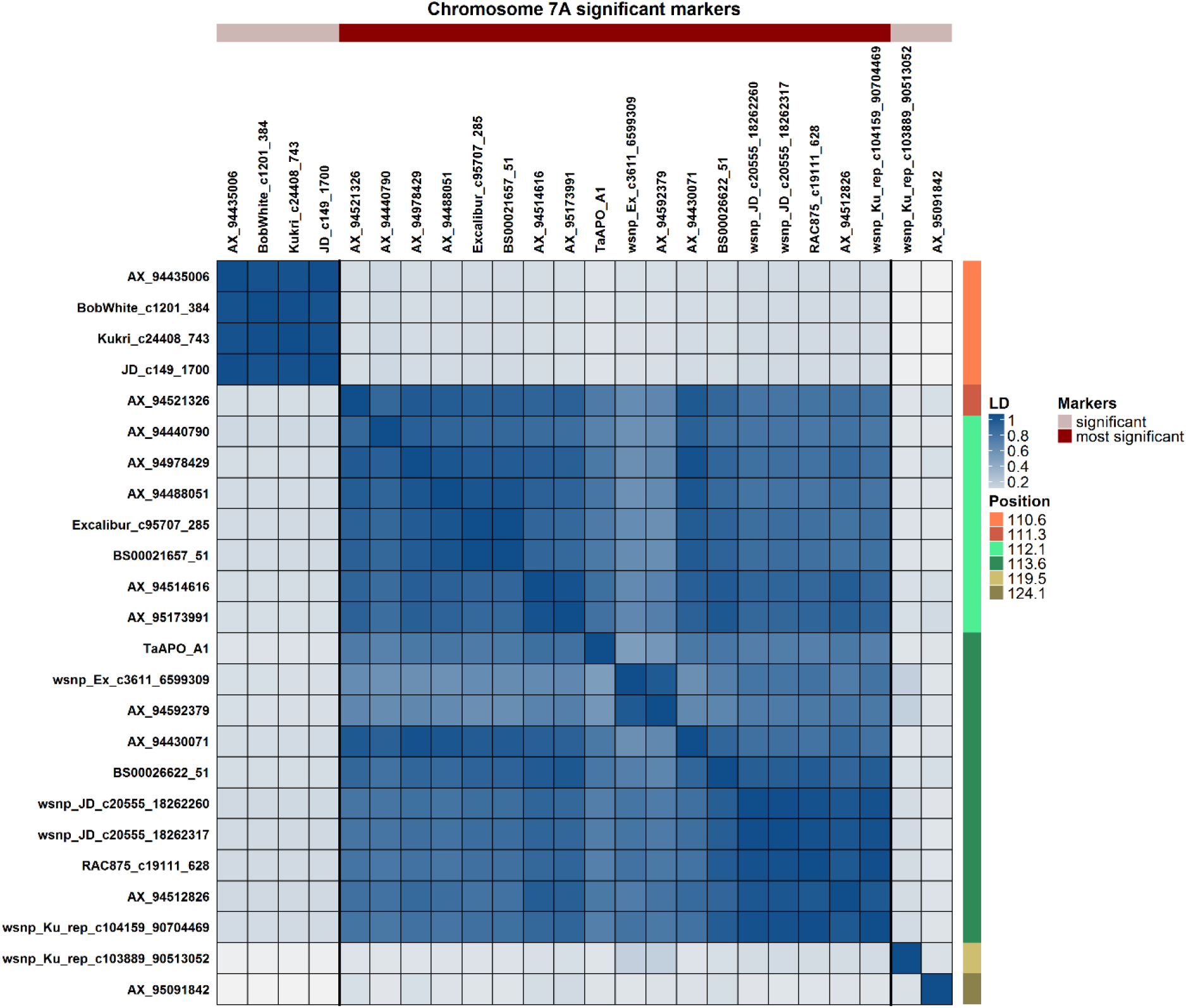
Pairwise linkage disequilibrium (*r*^2^) among the marker loci (including the KASP marker for *TaAPO-A1*) present in significant genetic region of TSN on chromosome 7A in wheat. Based on the linkage blocks, markers are divided into two categories viz., significant, and most significant. The color key is given in the figure.

The single nucleotide substitution G (low TSN allele) to T (high TSN allele) in the conserved functional domain of TaAPO-A1 resulted in a non-synonymous amino acid substitution from cysteine (C) to phenylalanine (F). The amino acid cysteine appears to be well conserved across various grass species at this position potentially indicating the conservation of C residue across grasses. However, the SIFT (Sorting Intolerant from Tolerant) score (Sim et al. 2012) analysis showed no potential deleterious effect from C to F substitution at this position (Table S3). We then looked at the promoter region of *TaAPO-A1* in ten genotypes from *The 10+ Wheat Genome Project* and identified a 115 bp INDEL (insertion-deletion) polymorphism at −484 bp upstream of the transcription start site of *TaAPO-A1*. Interestingly, the low TSN haplotype “G” (coding for cysteine) always had a deletion of 115 bp in the promoter, whereas the high TSN haplotype “T” (coding for phenylalanine) had 115 bp insertion. It, nevertheless, remains to be established via functional studies if this INDEL affects the transcription rate of *TaAPO-A1* contributing to the observed phenotypic differences for TSN in two haplogroups.

### Phylogenetic analyses show that TaAPO-A1, an ortholog of UFO in Arabidopsis, is conserved across terrestrial plant species

The BLAST search of *TaAPO-A1* orthologs across diverse plant species from the EnsemblPlants and the protein databases Phytozome v12.1 retrieved 64 protein sequences from 37 genera (52 species, Table S4) including Bryophytes, eudicots, and monocots. The final alignment consisted of 670 positions. The obtained ML topology reflects the evolution of terrestrial plants with *Amborella trichopoda* at the basis of the two main clades, monocotyledons and eudicotyledons (Fig. 6). The protein is relatively well conserved as seen from the very small branches especially within the grass tribe Triticeae, including *Triticum aestivum* and *Hordeum vulgare*, which diverged about ten million years ago (Ma) (Bernhardt et al. 2017) or even the Poaceae, whose most recent common ancestor probably occurred 50-75 Ma (Bouchenak-Khelladi et al. 2010).

**Figure 6:**
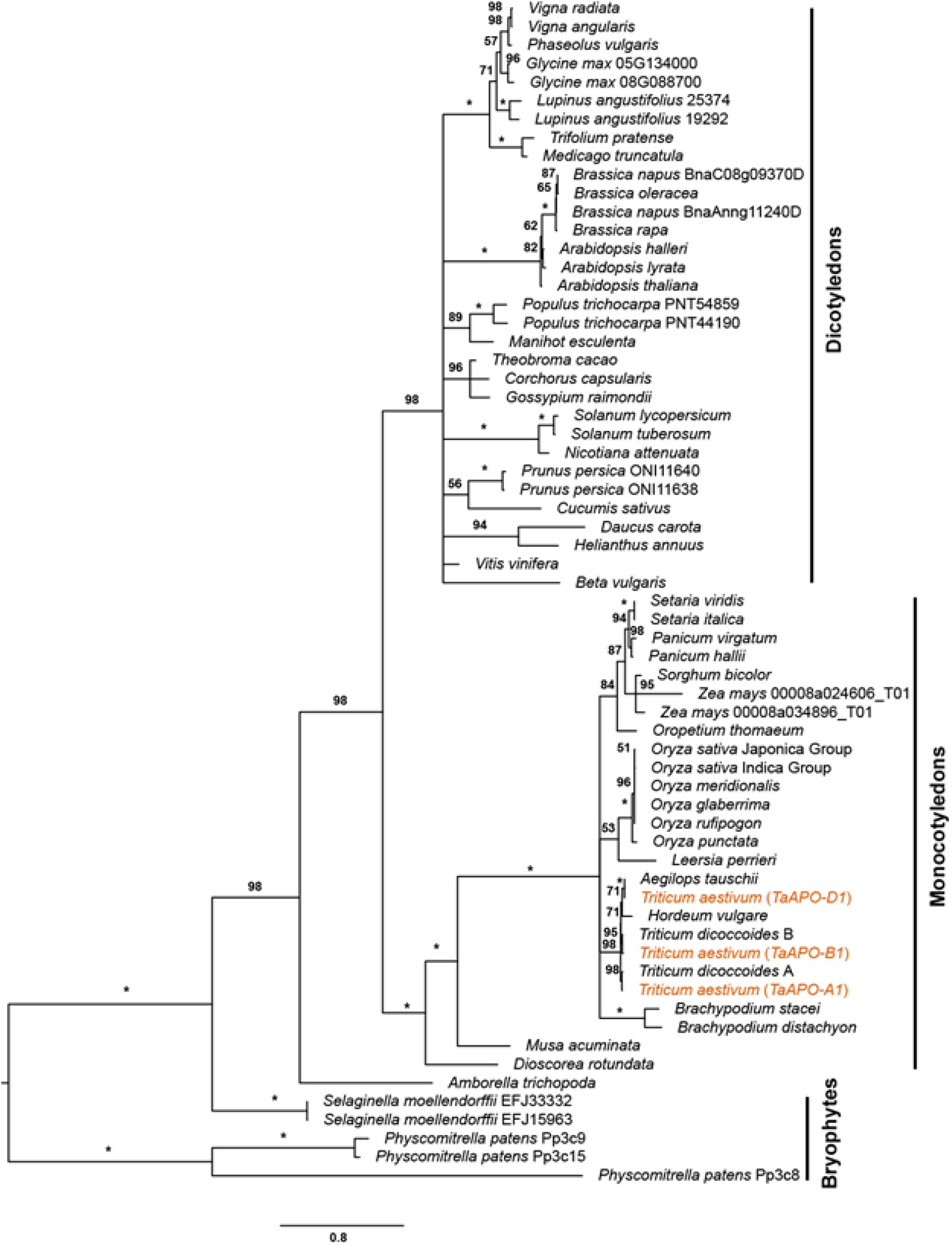
Maximum likelihood phylogenetic tree of *TaAPO-A1* orthologous proteins across terrestrial plant species. Bootstrap values are indicated along the braches. Asterisks indicate >99% bootstrap values. The *TaAPO* homoeologs are highlighted in orange color. The bars on the right side indicate the major clades. The amino acid substitution scale is indicated at the bottom of the figure.

## Discussion

### Exploiting significant, heritable genetic variation of TSN as well as a positive correlation with other traits can help to improve the grain yield in wheat

Grain yield (GY) improvement is considered as the top focus of virtually every wheat breeding program. However, an extremely complex genetic nature of GY often hampers its genetic improvement as it is the product of several yield components, e.g., the number of spikes per plant, grains per spike, thousand-grain weight. The number of grains per spike is a product of TSN and fertility. Therefore, an essential consideration in wheat breeding has been to employ a reductionist approach, i.e., to exploit the information about the individual component traits; most of which are negatively associated with each other. In this study, we analyzed a winter wheat panel comprising of 518 varieties for grain yield component traits such as TSN and SL along with the flowering time (FT). The grain yield data based on previous studies were taken for comparison purposes (Schulthess et al. 2017). In all observed traits, besides significant genetic variation, we observed a significant genotype-by-environment (year) interaction. Nevertheless, the broad-sense heritability estimates ranging from 0.68 to 0.89 suggested that genetic variation is heritable – an essential indicator of high selection response (Table 1). Similar heritability values for the studied traits have been reported recently in other diverse mapping populations (Guo et al. 2017; Würschum et al. 2018).

In addition to significant genetic variation, TSN showed a positive and significant correlation with SL, FT, and GY (Fig. 1e). This showed that albeit being weak (which is by virtue of the extreme quantitative genetic nature of GY), the correlation with GY could help improve the genetic gain. Moreover, it should be noted that the genetic architecture of yield component traits *per se* is also important which means that if the component traits possess complex genetic architecture, the problem of grain yield improvement would be further compounded. Nevertheless, a reasonably high heritability value suggests that TSN is strongly genetically inherited and that mapping of the underlying quantitative trait loci (QTL) would be efficient.

### High marker density governs the efficacy of genetic and physical mapping

The efficiency of GWAS depends on the size of the population and genetic diversity. Genome-wide marker density with many polymorphic sites is therefore vital and coupled with a sharp decline in linkage disequilibrium (LD) between marker loci; it increases GWAS resolution. In our study, the size of the population, high-density genotyping, and the use of stringent linear mixed-effect models warranted the genetic mapping of true marker-trait-associations (MTA). As noted in another study based on a subset of varieties, the absence of distinct sub-populations in this panel suggests that the European winter wheat varieties have been bred, by and large, from a narrow genetic base and with similar goals (Muqaddasi et al. 2019) which is in line with other reports based on studies using similar genetic material but different marker platforms (Kollers et al. 2013; Würschum et al. 2013).

To identify the candidate genes, high marker density in a given QTL genetic region is necessary since it helps to narrow-down to the physical region harboring the gene underlying the trait. Moreover, since GWAS hinges on the principle that markers work as proxies to the genes/QTL underlying the traits, a high density of markers in the QTL genetic region becomes vital for the success of fine mapping. In this study, we exploited this premise to identify a candidate gene physically.

### Physical mapping shows that TaAPO-A1 is a likely candidate gene for TSN in wheat

Our GWAS analysis revealed a significant QTL for total spikelet number on chromosome 7A, which explained ~25% of the total genotypic variance. Also, Würschum et al. (2018) recently reported a QTL for TSN on chromosome 7A in a similar type of elite winter wheat germplasm. Zhang et al. (2015) reported a putative *MOC1* ortholog to be associated with spikelet number, which is also located on chromosome 7A.

The strategy to investigate orthologous genes of rice with the known function was already successfully applied for various genes associated with grain size, grain weight as well as yield in wheat (Ma et al. 2016; Su et al. 2011; Wang et al. 2015; Zhang et al. 2012; Zhang et al. 2014; Zheng et al. 2014). The highly significant region of the detected TSN-QTL in our study corresponded to a physical interval of <1 Mb, containing a block of only ten genes, all in high LD (Figure 5). Based on the functional annotations, the rice gene *ABERRANT PANICLE ORGANIZATION 1* (*APO1*), an ortholog of *Arabidopsis UFO* (Ikeda et al. 2007; Ikeda et al. 2005; Samach et al. 1999; Wilkinson and Haughn 1995) was considered as the most likely candidate gene and was named as *TaAPO-A1* in wheat. The functional analyses in both rice and *Arabidopsis* revealed that the F-box containing protein is involved in the regulation and development of floral organs; more specifically *APO1* in rice that controls the number of spikelets per panicle by regulating the cell proliferation in meristems (Ikeda-Kawakatsu et al. 2009).

### Functional diversity among orthologs of TaAPO-A1 reveals the conserved F-box domain

The availability of genomic data for several wheat varieties from *The 10+ Wheat Genome Project* allowed the investigation of the intraspecific diversity of *TaAPO-A1* gene among wheat varieties. The *TaAPO-A1* contains two exons, each containing a SNP which causes an amino acid substitution. In the first exon, a T/G polymorphism at base 140 was related to the exchange of phenylalanine to cysteine, and in the second exon, at base 1284, a G/A polymorphism mutated aspartic acid to asparagine (Figure S6). It was possible to develop a functional KASP marker for the SNP in the first exon and to screen the germplasm panel. Both alleles were present in almost identical frequencies with 49.38% of the varieties carrying the allele of Chinese Spring with nucleotide T (referred to as *TaAPO-A1*b) and 50.62% of the varieties carrying the G nucleotide (referred to as *TaAPO-A1*a). The Chinese Spring allele was strongly associated (*P* <2.2e-16) with an increase in TSN and moderately associated with an increase in spike length (*P* = 3.4e-04) and yield (*P* = 9.0e-04) (Figure 4). For the B- and D-genomes, the orthologs of *TaAPO-A1* were related to the genes *TraesCS7B01G384000* and *TraesCS7D01G468700*. However, no MTAs were discovered on these genomes. The identified *TaAPO-A1* variants reflect natural allelic diversity with mild phenotypic effects, which is beneficial for practical breeding.

The presence of *TaAPO-A1* orthologs in a wide range of plants including Bryophytes, monocotyledons, and eudicotyledons suggests a central role of this gene class in the evolution and development of terrestrial plants (Figures 6, S7). The *Arabidopsis* gene *UFO* and rice *APO1* (orthologs of *TaAPO-A1*) encode for an F-box containing protein. It has been shown that the rice *APO1* and *Arabidopsis UFO* are important for floral development in respective species (Ikeda et al. 2007; Samach et al. 1999). Molecularly, the proteins SKP1, cullin like and F-box containing polypeptides form SCF protein complexes to function as E3-ubiquitin ligases that target specific proteins for degradation (Kaiser et al. 1998; Patton et al. 1998). It has been shown that *Arabidopsis UFO* indirectly regulates the expression of class B floral homeotic gene *APETALA 3* by targeting the degradation of proteins which negatively regulate its transcription (Samach et al. 1999). The rice *apo1* mutants show a reduction in the number of primary branches and thereby the number of spikelets due to the precocious conversion of inflorescence meristem (IM) to spikelet meristem (SM). Such a mutant phenotype offers an indication that *APO1* might target proteins that promote the precocious conversion of IM to SM for degradation in a functional state. In line with this idea, the dominant gain of function *APO1* alleles with an elevated expression as well as overexpression transgenic lines of *APO1* showed prolonged inflorescence development resulting in more branch iterations and consequently more spikelets (Ikeda-Kawakatsu et al. 2009).

From our promoter analysis, we found an INDEL where the 115 bp insertion was always associated with high TSN haplotype, whereas the deletion with low TSN haplotype. From this finding, it may be inferred that winter wheat genotypes in the haplogroup with insertion polymorphism have slightly elevated expression of *TaAPO- A1* leading to prolonged maturation of inflorescence meristem eventually producing more spikelets per spike. Conversely, the deletion haplotype has a comparatively reduced expression level of *TaAPO-A1*, leading to less number of spikelets. Nevertheless, validation of the INDEL haplotype across the whole winter wheat panel as well as expression analysis of *TaAPO-A1* in the two haplogroups with high and low TSN may offer further insights into the regulation of TSN in wheat.

## Conclusions

Our results demonstrate that with the availability of modern genomic tools such as the wheat reference sequence and the access to *The 10+ Wheat Genome Project*, the way from phenotype to a candidate gene is shortened considerably. Nevertheless, a robust genetic analysis including appropriate mapping populations, accurate and high-density genotyping, and proper phenotypic analyses are prerequisites to detecting significant QTL regions from which the causative genes could be deduced.

## Acknowledgments

The genotyping data were produced in the project VALID funded by the German Federal Ministry of Education and Research (BMBF; project number 0315947). We are grateful to Ellen Weiß, Anette Heber, Ute Ostermann, and Sonja Allner for help in phenotypic data collection. We are thankful to *The 10+ Wheat Genome Project* for making the resources available before publication.

## Author contribution statement

QHM and MSR conceived the idea. QHM analyzed the data, interpreted the results, and wrote the manuscript. JB and RK contributed to sequence and phylogenetic analyses. JP and MWG contributed the genotypic data. RK and MSR contributed to the interpretation of results and writing of the manuscript.

## References

Allen AM, Winfield MO, Burridge AJ, Downie RC, Benbow HR, Barker GL, Wilkinson PA, Coghill J, Waterfall C, Davassi A, Scopes G, Pirani A, Webster T, Brew F, Bloor C, Griffiths S, Bentley AR, Alda M, Jack P, Phillips AL, Edwards KJ (2017) Characterization of a Wheat Breeders’ Array suitable for high-throughput SNP genotyping of global accessions of hexaploid bread wheat (*Triticum aestivum*). Plant Biotechnol J 15:390–401

Altschul SF, Gish W, Miller W, Myers EW, Lipman DJ (1990) Basic local alignment search tool. Journal of Molecular Biology 215:403–410

Benjamini Y, Hochberg Y (1995) Controlling the false discovery rate: a practical and powerful approach to multiple testing. Journal of the Royal Statistical Society Series B (Methodological):289–300

Bernhardt N, Brassac J, Kilian B, Blattner FR (2017) Dated tribe-wide whole chloroplast genome phylogeny indicates recurrent hybridizations within Triticeae. BMC Evolutionary Biology 17:141

Boden SA, Cavanagh C, Cullis BR, Ramm K, Greenwood J, Finnegan EJ, Trevaskis B, Swain SM (2015) *Ppd-1* is a key regulator of inflorescence architecture and paired spikelet development in wheat. Nature Plants 1:14016

Bouchenak-Khelladi Y, Verboom GA, Savolainen V, Hodkinson TR (2010) Biogeography of the grasses (Poaceae): a phylogenetic approach to reveal evolutionary history in geographical space and geological time. Botanical Journal of the Linnean Society 162:543–557

Consortium IWGS (2018) Shifting the limits in wheat research and breeding using a fully annotated reference genome. Science 361:eaar7191

Darriba D, Taboada GL, Doallo R, Posada D (2011) ProtTest 3: fast selection of bestfit models of protein evolution. Bioinformatics 27:1164–1165

Debernardi JM, Lin H, Chuck G, Faris JD, Dubcovsky J (2017) microRNA172 plays a crucial role in wheat spike morphogenesis and grain threshability. Development 144:1966–1975

Deng Z, Cui Y, Han Q, Fang W, Li J, Tian J (2017) Discovery of consistent QTLs of wheat spike-related traits under nitrogen treatment at different development stages. Front Plant Sci 8:2120

Faris JD, Fellers JP, Brooks SA, Gill BS (2003) A bacterial artificial chromosome contig spanning the major domestication locus Q in wheat and identification of a candidate gene. Genetics 164:311–321

Frichot E, François O (2015) LEA: An R package for landscape and ecological association studies. Methods in Ecology and Evolution 6:925–929

Gauley A, Boden SA (2019) Genetic pathways controlling inflorescence architecture and development in wheat and barley. Journal of Integrative Plant Biology 61:296–309

Greenwood JR, Finnegan EJ, Watanabe N, Trevaskis B, Swain SM (2017) New alleles of the wheat domestication gene *Q* reveal multiple roles in growth and reproductive development. Development 144:1959–1965

Guindon S, Gascuel O (2003) A simple, fast, and accurate algorithm to estimate large phylogenies by maximum likelihood. Systematic Biology 52:696–704

Guo Z, Chen D, Alqudah AM, Röder MS, Ganal MW, Schnurbusch T (2017) Genome-wide association analyses of 54 traits identified multiple loci for the determination of floret fertility in wheat. New Phytologist 214:257–270

Guo Z, Zhao Y, Röder MS, Reif JC, Ganal MW, Chen D, Schnurbusch T (2018) Manipulation and prediction of spike morphology traits for the improvement of grain yield in wheat. Sci Rep-Uk 8:14435

Ikeda-Kawakatsu K, Yasuno N, Oikawa T, Iida S, Nagato Y, Maekawa M, Kyozuka J (2009) Expression level of *ABERRANT PANICLE ORGANIZATION1* determines rice inflorescence form through control of cell proliferation in the meristem. Plant Physiol 150:736–747

Ikeda K, Ito M, Nagasawa N, Kyozuka J, Nagato Y (2007) Rice *ABERRANT PANICLE ORGANIZATION 1*, encoding an F-box protein, regulates meristem fate. The Plant Journal 51:1030–1040

Ikeda K, Nagasawa N, Nagato Y (2005) *ABERRANT PANICLE ORGANIZATION 1* temporally regulates meristem identity in rice. Developmental Biology 282:349–360

Jones DT, Taylor WR, Thornton JM (1992) The rapid generation of mutation data matrices from protein sequences. Bioinformatics 8:275–282

Kaiser P, Sia RA, Bardes EG, Lew DJ, Reed SI (1998) Cdc34 and the F-box protein Met30 are required for degradation of the Cdk-inhibitory kinase Swe1. Genes & Development 12:2587–2597

Kearse M, Moir R, Wilson A, Stones-Havas S, Cheung M, Sturrock S, Buxton S, Cooper A, Markowitz S, Duran C (2012) Geneious Basic: an integrated and extendable desktop software platform for the organization and analysis of sequence data. Bioinformatics 28:1647–1649

Kollers S, Rodemann B, Ling J, Korzun V, Ebmeyer E, Argillier O, Hinze M, Plieske J, Kulosa D, Ganal MW, Röder MS (2013) Genetic architecture of resistance to Septoria tritici blotch (*Mycosphaerella graminicola*) in European winter wheat. Molecular Breeding 32:411–423

Koppolu R, Schnurbusch T (2019) Developmental pathways for shaping spike inflorescence architecture in barley and wheat. Journal of Integrative Plant Biology 61:278–295

Li C, Lin H, Chen A, Lau M, Jernstedt J, Dubcovsky J (2019) Wheat *VRN1* and *FUL2* play critical and redundant roles in spikelet meristem identity and spike determinacy. bioRxiv:510388

Liu J, Xu Z, Fan X, Zhou Q, Cao J, Wang F, Ji G, Yang L, Feng B, Wang T (2018) A Genome-Wide Association Study of Wheat Spike Related Traits in China. Front Plant Sci 9

Ma L, Li T, Hao C, Wang Y, Chen X, Zhang X (2016) *TaGS5-3A*, a grain size gene selected during wheat improvement for larger kernel and yield. Plant Biotechnol J 14:1269–1280

Muqaddasi QH, Zhao Y, Rodemann B, Plieske J, Ganal MW, Röder MS (2019) Genome-wide Association Mapping and Prediction of Adult Stage *Septoria tritici* Blotch Infection in European Winter Wheat via High-Density Marker Arrays. The Plant Genome 12:180029

Ochagavía H, Prieto P, Savin R, Griffiths S, Slafer G (2018) Dynamics of leaf and spikelet primordia initiation in wheat as affected by *Ppd-1a* alleles under field conditions. J Exp Bot 69:2621–2631

Patton EE, Willems AR, Tyers M (1998) Combinatorial control in ubiquitin-dependent proteolysis: don’t Skp the F-box hypothesis. Trends in Genetics 14:236–243

Sakuma S, Golan G, Guo Z, Ogawa T, Tagiri A, Sugimoto K, Bernhardt N, Brassac J, Mascher M, Hensel G, Ohnishi S, Jinno H, Yamashita Y, Ayalon I, Peleg Z, Schnurbusch T, Komatsuda T (2019) Unleashing floret fertility in wheat through the mutation of a homeobox gene. Proceedings of the National Academy of Sciences 116:5182–5187

Samach A, Klenz JE, Kohalmi SE, Risseeuw E, Haughn GW, Crosby WL (1999) The *UNUSUAL FLORAL ORGANS* gene of *Arabidopsis thaliana* is an F-box protein required for normal patterning and growth in the floral meristem. The Plant Journal 20:433–445

Schulthess AW, Reif JC, Ling J, Plieske J, Kollers S, Ebmeyer E, Korzun V, Argillier O, Stiewe G, Ganal MW, Röder MS, Jiang Y (2017) The roles of pleiotropy and close linkage as revealed by association mapping of yield and correlated traits of wheat (*Triticum aestivum* L.). J Exp Bot 68:4089–4101

Shaw LM, Lyu B, Turner R, Li C, Chen F, Han X, Fu D, Dubcovsky J (2018) *FLOWERING LOCUS T2* regulates spike development and fertility in temperate cereals. J Exp Bot 70:193–204

Sim N-L, Kumar P, Hu J, Henikoff S, Schneider G, Ng PC (2012) SIFT web server: predicting effects of amino acid substitutions on proteins. Nucleic acids research 40:W452–W457

Sorrells ME, Gustafson JP, Somers D, Chao S, Benscher D, Guedira-Brown G, Huttner E, Kilian A, McGuire PE, Ross K (2011) Reconstruction of the Synthetic W7984 × Opata M85 wheat reference population. Genome 54:875–882

Stamatakis A (2014) RAxML version 8: a tool for phylogenetic analysis and post-analysis of large phylogenies. Bioinformatics 30:1312–1313

Su Z, Hao C, Wang L, Dong Y, Zhang X (2011) Identification and development of a functional marker of *TaGW2* associated with grain weight in bread wheat (*Triticum aestivum* L.). Theor Appl Genet 122:211–223

Utz HF, Melchinger AE, Schön CC (2000) Bias and sampling error of the estimated proportion of genotypic variance explained by quantitative trait loci determined from experimental data in maize using cross validation and validation with independent samples. Genetics 154:1839–1849

VanRaden PM (2008) Efficient methods to compute genomic predictions. Journal of Dairy Science 91:4414–4423

Wang S, Zhang X, Chen F, Cui D (2015) A single-nucleotide polymorphism of *TaGS5* gene revealed its association with kernel weight in Chinese bread wheat. Front Plant Sci 6:1166

Wang SC, Wong DB, Forrest K, Allen A, Chao SM, Huang BE, Maccaferri M, Salvi S, Milner SG, Cattivelli L, Mastrangelo AM, Whan A, Stephen S, Barker G, Wieseke R, Plieske J, International Wheat Genome Sequencing Consortium, Lillemo M, Mather D, Appels R, Dolferus R, Brown-Guedira G, Korol A, Akhunova AR, Feuillet C, Salse J, Morgante M, Pozniak C, Luo MC, Dvorak J, Morell M, Dubcovsky J, Ganal M, Tuberosa R, Lawley C, Mikoulitch I, Cavanagh C, Edwards KJ, Hayden M, Akhunov E (2014) Characterization of polyploid wheat genomic diversity using a high-density 90 000 single nucleotide polymorphism array. Plant Biotechnol J 12:787–796

Wilkinson MD, Haughn GW (1995) *UNUSUAL FLORAL ORGANS* controls meristem identity and organ primordia fate in Arabidopsis. The Plant Cell 7:1485–1499

Würschum T, Langer SM, Longin CFH, Korzun V, Akhunov E, Ebmeyer E, Schachschneider R, Schacht J, Kazman E, Reif JC (2013) Population structure, genetic diversity and linkage disequilibrium in elite winter wheat assessed with SNP and SSR markers. Theor Appl Genet 126:1477–1486

Würschum T, Leiser WL, Langer SM, Tucker MR, Longin CFH (2018) Phenotypic and genetic analysis of spike and kernel characteristics in wheat reveals long-term genetic trends of grain yield components. Theor Appl Genet 131:2071–2084

Yu JM, Pressoir G, Briggs WH, Bi IV, Yamasaki M, Doebley JF, McMullen MD, Gaut BS, Nielsen DM, Holland JB, Kresovich S, Buckler ES (2006) A unified mixed-model method for association mapping that accounts for multiple levels of relatedness. Nat Genet 38:203–208

Zhai H, Feng Z, Li J, Liu X, Xiao S, Ni Z, Sun Q (2016) QTL analysis of spike morphological traits and plant height in winter wheat (*Triticum aestivum* L.) using a high-density SNP and SSR-based linkage map. Front Plant Sci 7:1617

Zhang B, Liu X, Xu W, Chang J, Li A, Mao X, Zhang X, Jing R (2015) Novel function of a putative *MOC1* ortholog associated with spikelet number per spike in common wheat. Sci Rep-Uk 5:12211

Zhang L, Zhao YL, Gao LF, Zhao GY, Zhou RH, Zhang BS, Jia JZ (2012) *TaCKX6-D1*, the ortholog of rice *OsCKX2*, is associated with grain weight in hexaploid wheat. New Phytologist 195:574–584

Zhang Y, Liu J, Xia X, He Z (2014) *TaGS-D1*, an ortholog of rice *OsGS3*, is associated with grain weight and grain length in common wheat. Molecular breeding 34:1097–1107

Zheng J, Liu H, Wang Y, Wang L, Chang X, Jing R, Hao C, Zhang X (2014) *TEF-7A*, a transcript elongation factor gene, influences yield-related traits in bread wheat (*Triticum aestivum* L.). J Exp Bot 65:5351–5365

