## Supplementary Figs. 1-5 for "*TaAPO-A1*, an ortholog of rice *ABERRANT PANICLE ORGANIZATION 1*, is associated with total spikelet number per spike in elite hexaploid winter wheat varieties (*Triticum aestivum* L.)"

Article type: Research article

### Supplementary figures

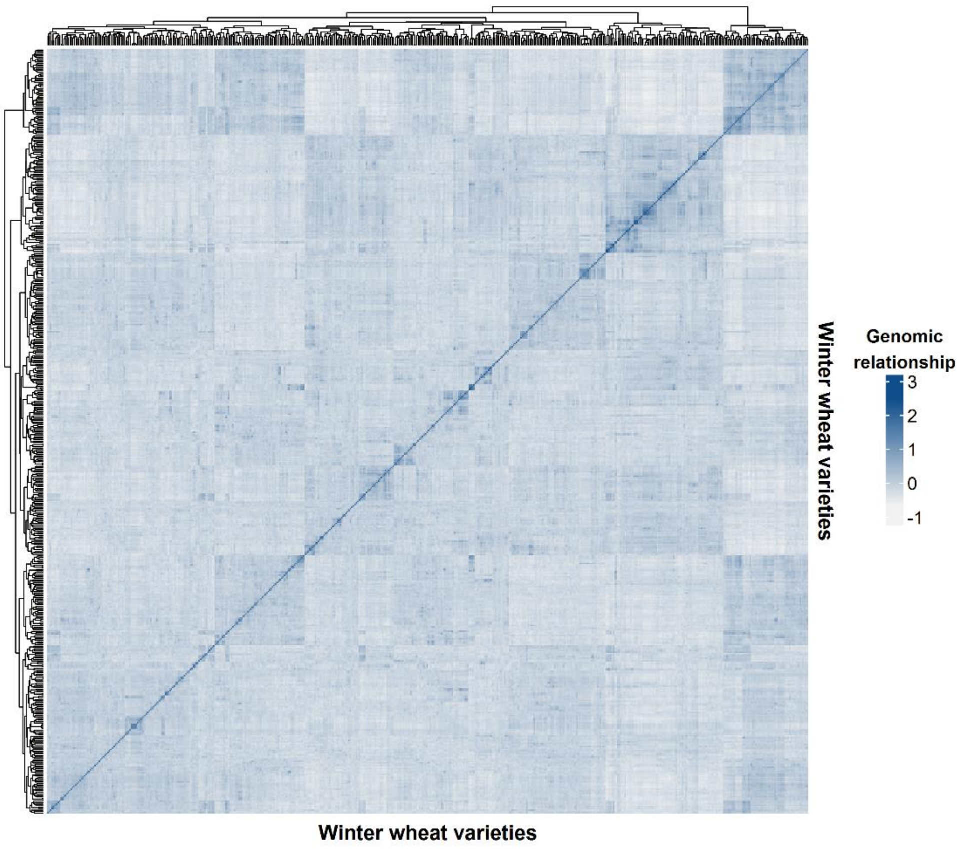


**Figure S1. Heat map of the genomic relationships among 518 European winter wheat varieties based on 39,908 marker genotypes.** The varieties are ordered according to hierarchical cluster analysis.

**(a)**

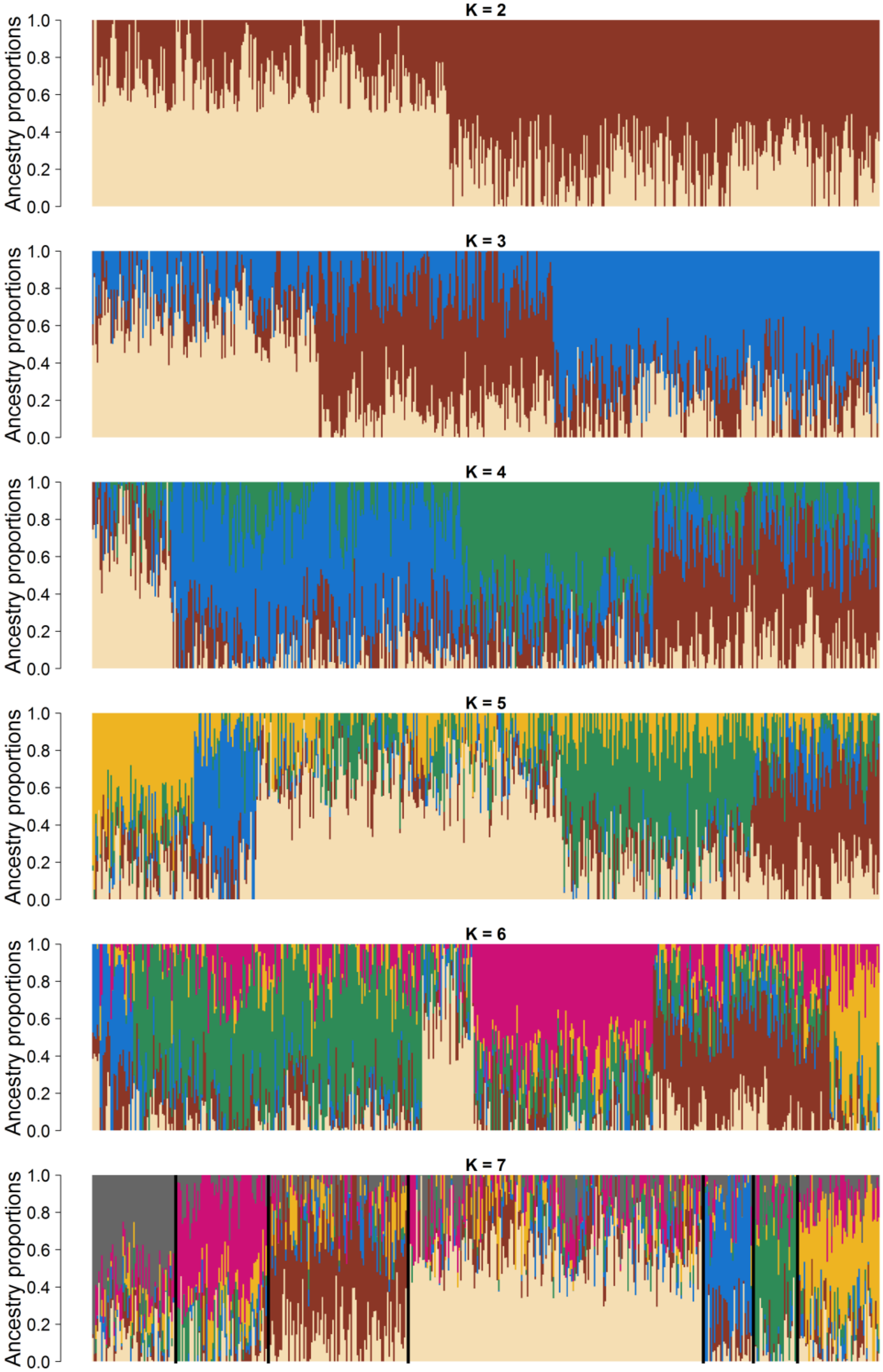

**(b)**

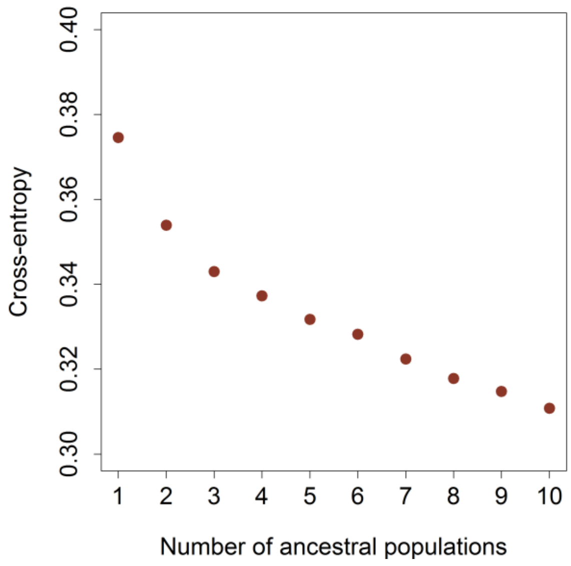


**Figure S2. Population structure analysis of 518 European winter wheat varieties based on 39,809 marker genotypes. (a) Bar plots show the existence of admixed sub-populations, (b) The cross-entropy plot shows that there exists a minimal sub-structuring in the panel.**

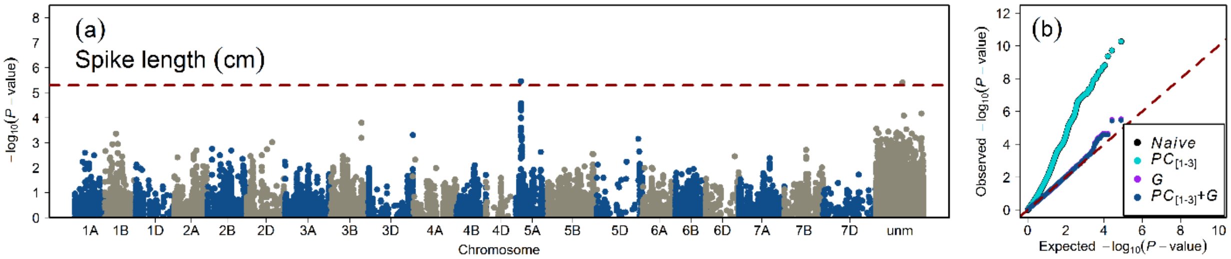


**Figure S3: Summary of genome-wide association studies of spike length in the population of 518 European winter wheat varieties. (a) Manhattan plot shows the distribution of marker significance $-\log_{10}(P - \text{value})$ along the chromosomes.** The correction for population stratification and familial relatedness was performed by using the first three principal components (*PC*[1-3]) and an additive genomic relationship matrix (*G*) in a linear mixed-effect model. The red dashed line marks the multiple testing criteria of false discovery rate (FDR) <0.10, **(b) Quantile-quantile plot showing the distribution of observed versus expected (red dashed line) $-\log_{10}(P - \text{value})$.** The general linear model (*naive*) without correction for population structure, the *PC*[1-3] model (population structure corrected with the first three *PCs*), the *G* model (familial relatedness corrected with a genomic relationship matrix), and the *PC*[1-3]+*G* model (population structure and familial relatedness corrected with *PCs* and the *G* matrix). The color code for different models is given in the figure legend.

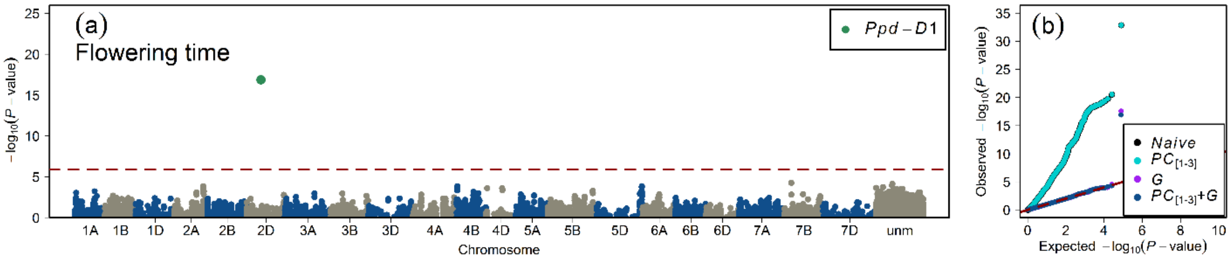


**Figure S4: Summary of genome-wide association studies of flowering time in the population of 518 European winter wheat varieties. (a) Manhattan plot shows the distribution of marker significance $-\log_{10}(P - \text{value})$ along the chromosomes.** The correction for population stratification and familial relatedness was performed by using the first three principal components (*PC*[1-3]) and an additive genomic relationship matrix (*G*) in a linear mixed-effect model. The red dashed line marks the multiple testing criteria of false discovery rate (FDR) <0.05, **(b) Quantile-quantile plot showing the distribution of observed versus expected (red dashed line) $-\log_{10}(P - \text{value})$.** The general linear model (*naive*) without correction for population structure, the *PC*[1-3] model (population structure corrected with the first three *PCs*), the *G* model (familial relatedness corrected with a genomic relationship matrix), and the *PC*[1-3]+*G* model (population structure and familial relatedness corrected with *PCs* and the *G* matrix). The color code for different models is given in the figure legend.

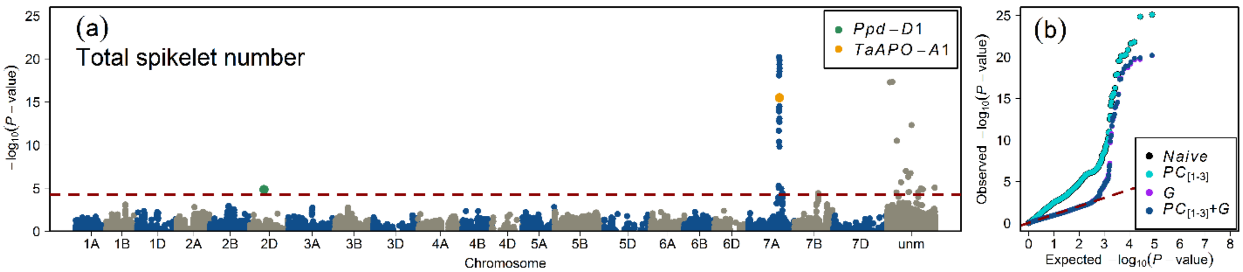


**Figure S5: Summary of genome-wide association studies of total spikelet number per spike in the population of 518 European winter wheat varieties. (a) Manhattan plot shows the distribution of marker significance $-\log_{10}(P-\text{value})$ along the chromosomes.** The correction for population stratification and familial relatedness was performed by using the first three principal components ($PC_{[1\text{-}3]}$) and an additive genomic relationship matrix (*G*) in a linear mixed-effect model. The red dashed line marks the multiple testing criteria of false discovery rate (FDR) <0.05, **(b) Quantile-quantile plot showing the distribution of observed versus expected (red dashed line) $-\log_{10}(P-\text{value})$.** The general linear model (*naive*) without correction for population structure, the $PC_{[1\text{-}3]}$ model (population structure corrected with the first three *PCs*), the *G* model (familial relatedness corrected with a genomic relationship matrix), and the $PC_{[1\text{-}3]}+G$ model (population structure and familial relatedness corrected with *PCs* and the *G* matrix). The color code for different models is given in the figure legend.
