## Supplementary figures and images for "*TaAPO-A1*, an ortholog of rice *ABERRANT PANICLE ORGANIZATION 1*, is associated with total spikelet number per spike in elite hexaploid winter wheat varieties (*Triticum aestivum* L.)"

### Supplementary Fig. 6

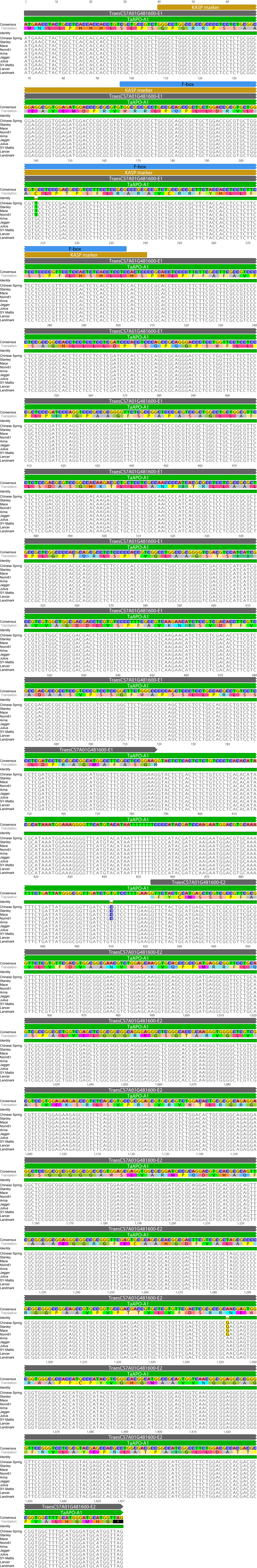

### Supplementary Fig. 7

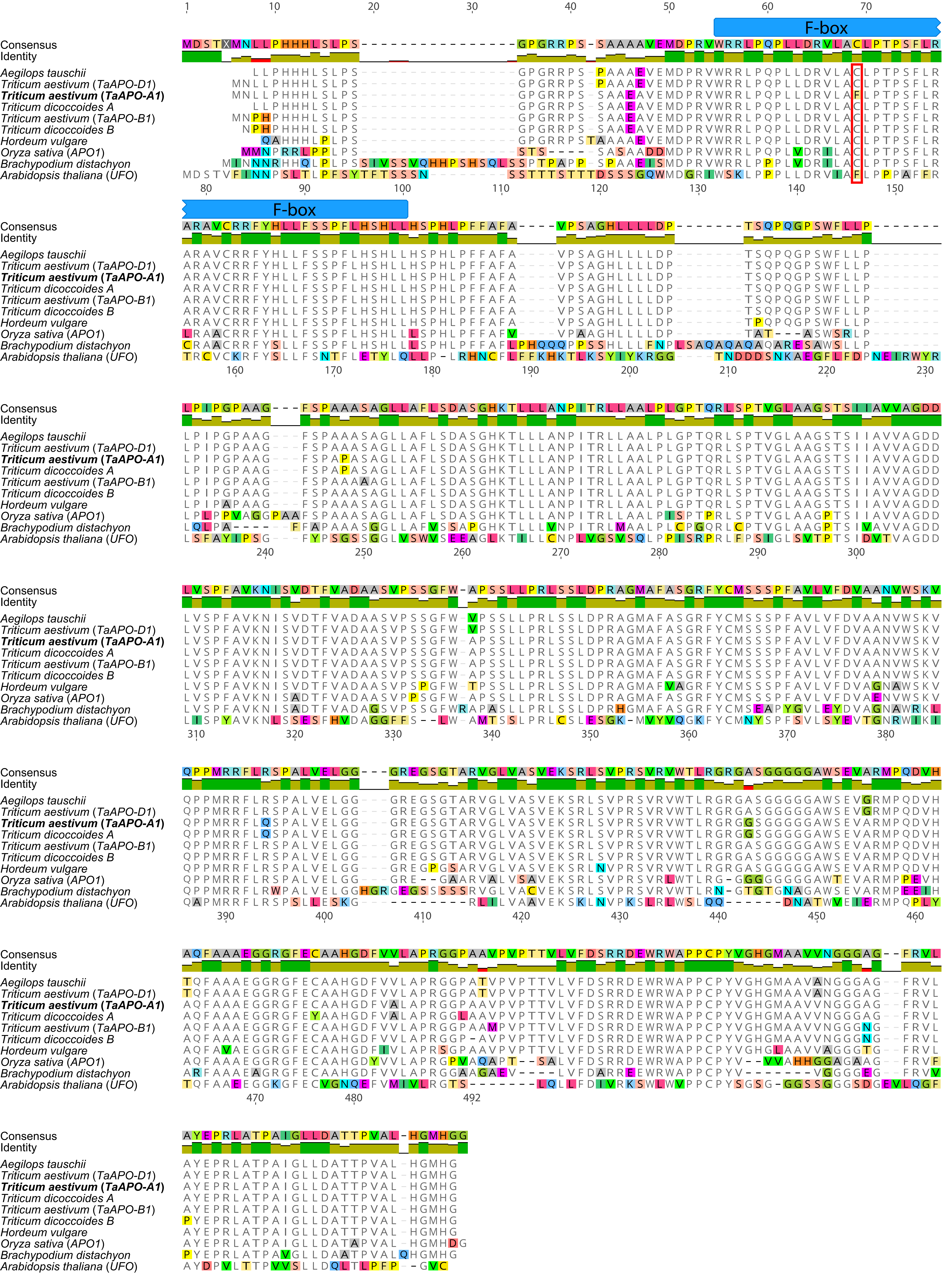
